# Dynamics Affecting the Risk of Silent Circulation When Oral Polio Vaccination Is Stopped

**DOI:** 10.1101/058099

**Authors:** J.S. Koopman, C.J Henry, J.H. Park, M.C. Eisenberg, E.L. Ionides, J.N. Eisenberg

## Abstract

Silent circulation of polioviruses without poliomyelitis cases could threaten eradication when oral polio vaccine (OPV) use is stopped worldwide. Waning immunity promotes silent circulation by increasing poliovirus transmission from individuals not at risk of paralytic polio. There is limited data on temporal patterns of waning. Accordingly, we modeled a range of waning patterns, scaled from fast but shallow to slow but deep, while keeping constant the effect of waning on transmission dynamics before vaccination begins. Besides waning, we varied overall transmissibility, the delay from beginning vaccination to reaching specified infection levels, and type specific virus characteristics. We observed an increasing range of vaccination levels that resulted in long periods of silent circulation after eliminating paralytic polio cases as the delay in reaching final vaccination levels increased. The extent of silent circulation was higher when waning was slower and deeper, when transmissibility was higher, and when virus was type 3. In our model, modest levels of vaccination of adults reduce silent circulation risks. These modeled patterns are consistent with very long silent circulation mainly emerging as a threat to OPV cessation in the last places from which polio cases are eliminated. Our analyses indicate why previous modeling studies have not seen the threat of silent circulation. They used models with no or very short duration waning and they lacked identifiability of waning effects on silent circulation because they fit models only to paralytic polio case counts. Our analyses show that nearly identical polio case count patterns can be generated by a range of waning patterns that in turn generate diverse silent circulation risks. We conclude that the risks of prolonged silent circulation are real but unquantified, that vaccinating adults with waned immunity will reduce those risks, and that intensive environmental surveillance will be needed to detect this risk before stopping OPV.

## Introduction

The final stages of polio eradication are approaching, after a campaign lasting almost 30 years that has encountered and overcome many foreseen and unforeseen challenges. Africa has now not seen a WPV caused polio case for more than a year. After high numbers of cases in Pakistan-Afghanistan in 2014, improved control programs lowered those numbers in 2015 and the expectation is there will be no new paralytic cases after 2016.

Focusing on paralytic cases detected by acute flaccid paralysis (AFP) surveillance (Grassly, 2013) has been sufficient to direct vaccination efforts where additional resources and improved methods are most needed to eliminate the last polio cases. Paralytic poliomyelitis cases occur almost exclusively after first infections. Transmissions between people who have been either previously infected or successfully vaccinated are silent in the sense that they are not detected by AFP surveillance. Ignoring such transmission events has not to date been an obstacle to eradicating polio from much of the world. Largely because of the success in focusing on primary infection, little is known about how much spread of virus occurs from reinfections of older individuals with waned immunity. In this paper we present analysis of reinfection dynamics that show how silent circulation fed by reinfections could present an emerging new challenge after elimination of polio cases in the West Africa and Pakistan-Afghanistan foci of transmission even though such silent circulation might not have created problems elsewhere. Silent circulation means transmission in the absence of polio cases.

The potential for prolonged silent circulation is important because the final polio eradication plan requires cessation of all oral polio vaccine (OPV) use (WHO, 2015). The reason for this decision is that the vaccine strain can cause paralysis and can be transmitted and evolve to cause paralysis and to transmit as readily as wild polio viruses (WPV) (Famulare et al., 2015).

Israel recently experienced more than a year of silent circulation. The virus that came to Israel had circulated silently from Pakistan into Egypt and then into Israel. The transmissions carrying virus across these long distances seem more likely to involve older individuals experiencing reinfections. But within Israel, administering OPV only to children who had received only IPV successfully eliminated virus transmission without a single paralytic case (Manor et al., 2014; Shulman et al., 2014a; Shulman et al., 2014b; Shulman et al., 2014c). From Israel the virus spread into Syria and Iraq where it caused an outbreak of paralytic cases (Arie, 2014; Aylward and Alwan, 2014). Other evidence that the potential for silent circulation is increasing includes increasing identification of orphan viruses not closely related to detected paralytic poliomyelitis cases (Hagan et al., 2015; Porter et al., 2015) despite improvements in AFP surveillance.

The potential for silent circulation to cause problems after the cessation of OPV use is now being experienced in Nigeria after the switch from trivalent to bivalent OPV stopped the use of OPV2. An environmental sample in Borno state detected a cVDPV2 whose sequence revealed it had evolved more than three years earlier and that had last caused a paralytic polio cases about two years earlier (GPEI, 2016).

Current thinking is that three years without paralytic poliomyelitis cases is sufficient to insure there is no ongoing circulation. This three year criterion is largely empirically based on experience in Latin America (Debanne and Rowland, 1998). It is theoretically supported initially using models that assumed acquired immunity never wanes (Eichner and Dietz, 1996; Kalkowska et al., 2012) and by a more recent paper assuming that significant waning only occurs in the first few years after infection (Kalkowska et al., 2015). A more recent analysis using a simple model with a hidden assumption that there was no long term waning also concluded the risks of silent circulation are ignorable after 3 years (Famulare, 2015). To date the models that incorporate waning immunity assume only fast and shallow waning, which is inconsistent with the observation of very deep levels of waning in elderly Dutch individuals (Abbink et al., 2005) and with recent vaccine trials whose control arms were consistent with considerable waning between the ages of 5 and 10 (Jafari et al., 2014). But those two studies used full dose vaccine exposures which might not reflect waning relevant to smaller, more realistic exposures.

To analyze the potential role of waning immunity in the continued transmission of polio after stopping OPV administration, we simplified a previous polio model we developed to examine success and failures of the polio eradication effort in the context of waning immunity and OPV transmissibility (Mayer et al., 2013). We converted the multiple step waning in that model to a single step to facilitate explorations of broad system dynamics and enable us to broadly sweep unknown parameters in ways that could direct more detailed models to productive explorations. Understanding those dynamics will help us understand the consequences of different eradication plans, determine which aspects of the unknown most urgently need to be studied, and build more informative models. Specifically, our analysis examines the impact of waning immunity dynamics on times to eliminate the last polio cases (i.e., sustained silent circulation) and how that might differ for different serotypes of polio. Our analysis seeks to uncover how and why the following interact to affect silent circulation risks: 1) waning immunity, 2) vaccination program deficiencies that cause them to take a long time to eliminate the last polio case, and 3) increasing transmission potential. It also seeks to develop theory to predict why different risks of silent circulation should be expected for different serotypes of polio and what actions before stopping OPV can help insure eradication.

## Methods

### Infection Transmission Model

We use a minimalist transmission model designed to capture the elements of the inference we seek while optimizing our capacity to clarify how those elements interact to produce prolonged low level silent circulation. More realistic and detailed models are available to capture the effects of waning and boosting of immunity (Behrend et al., 2014; Wagner et al., 2014). Those models, however, are fit to the high dose exposure of OPV administration. There is no data on which to formulate waning at low doses. Therefore we chose to examine a minimalist model to bring out joint effects between factors in a way that more complex models cannot, and to facilitate complex parameter sweeps and provide clarity on what is behind a given pattern of silent circulation in our model.

From the flow diagram in Figure 1 we see that our model distinguishes first infections from subsequent reinfections and WPV infections from OPV infections. Waning in the model occurs in a one-step transition from recovered state R to a partially susceptible state P. Recovery from all infections leads to a single immune state R. That means immunity from WPV or OPV infections is identical, as is immunity resulting from a first infection or a subsequent infection.

**Figure 1:**
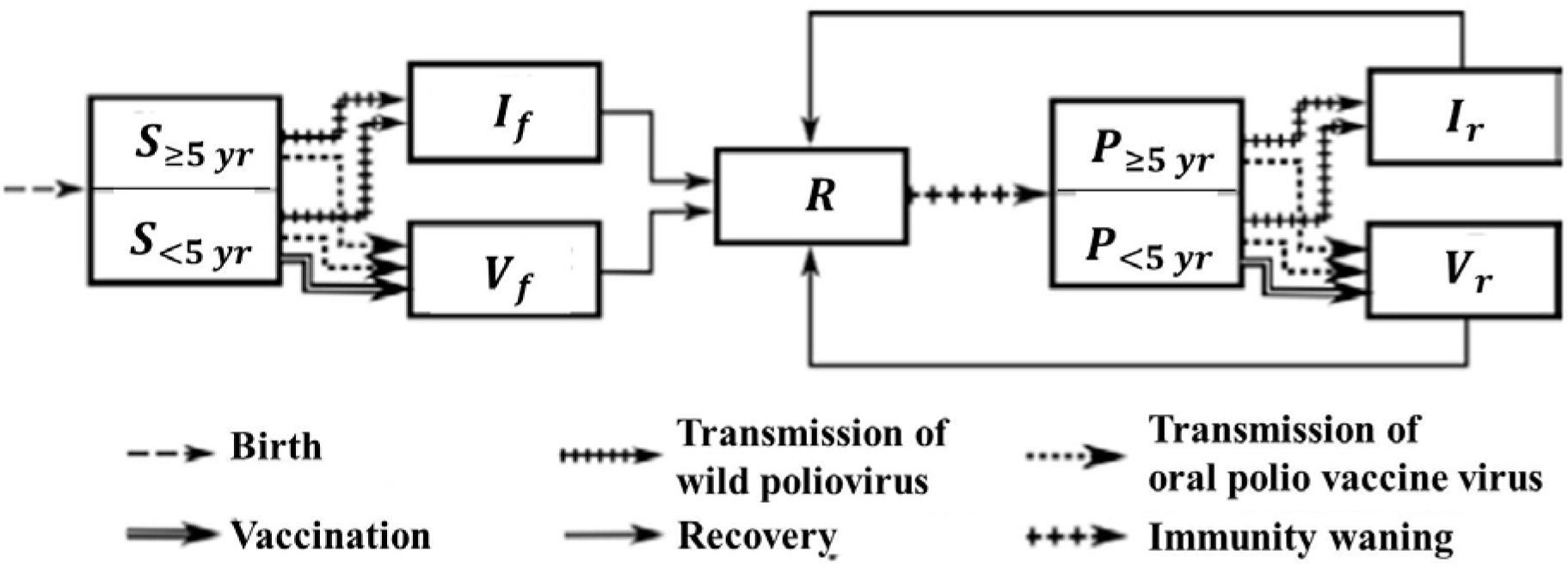
The seven compartments in our one step waning model and its flows not related to age or death are illustrated. The one waning step is from R to P. The speed of that flow is the speed of waning. The depth of waning is determined by the degree of susceptibility in the P compartment and the duration and contagiousness of the WPV or OPV infections that flow out of P. Although we distinguish between different age groups in all seven compartments (in order to allow for realistic aging), we only show here the distinctions that are relevant to infection processes, namely between (fully or partially) susceptible individuals who are or are not in the age group targeted by vaccination efforts.

#### More simplifying model assumptions

Our model analyzes only one polio serotype at a time. It uses parameters such as the fraction of first infections that result in paralytic poliomyelitis and the relative transmissibility of OPV compared to WPV, that have been used by others (Duintjer Tebbens et al., 2013c) to characterize the three serotypes of polio. It assumes homogeneous mixing and homogeneous contact rates across all compartments with paralytic cases transmitting identically to non-paralytic first infection cases. It assumes that death rates are constant, not affected by age, and are equal to birth rates. The differential equation model form used essentially assumes a continuous population rather than discrete individuals. It also assumes everyone in the population has an equal rate of contacting everyone else. Model equations and parameters are presented in the appendix.

The model does not directly distinguish paralytic infections from silent infections. But our analyses of model output use the first Infections to Paralytic infections Ratio (IPR). When the number of first infections in the past year first become less than the IPR, the time of the last paralytic polio case is declared. A new count of first infections begins at this time and goes on without time limit. When that total count exceeds the IPR, a time of reappearance or recurrence of polio cases declared. Our model assumes that reinfections do not cause paralytic poliomyelitis.

Eradication is said to occur in our model when the prevalence of infection falls below 1 in a population of one million. We explored a variety of definitions that counted different entities such as numbers of first and repeat infections weighted by their overall transmission potential. We found that none of these affected the basic conclusions of this paper.

In the real world, chance and population structure affect when the last paralytic polio case appears, when after that any new cases might appear, and when eradication occurs. By avoiding these strong realistic determinants we more clearly reveal the dynamics leading to prolonged silent circulation.

Different duration and contagiousness of infection are assigned to wild polio virus (WPV) and oral polio vaccine virus (OPV) infections. We assume that OPV infections have the same duration and contagiousness whether they arose from direct vaccination or transmission from someone with an OPV infection. To further reduce parameters we assume that ratios affecting relationships between first and reinfections are the same for OPV and WPV infections and that the ratios affecting duration and contagiousness of OPV compared to WPV infections are the same for first infections and reinfections.

#### Age structure

To examine vaccination effects we add age to the model because vaccination is only given to children less than 5 years old. Instead of making chronological time and aging time move across the same small time steps, we reduce the computations needed for model analysis by moving age progression across larger steps (1.5 months) than chronological time and keeping track of age only until age 5. The choice of aging step size resulted from a compromise between achieving sufficient speed of computations and avoiding an excessive discrepancy between the fraction of children who reach 5 years old without ever being vaccinated in our model and the corresponding fraction that would do so in the limiting case of infinitely small age steps.

#### Modeling vaccination patterns over time

Vaccination patterns over time were simplified by assuming that all children were vaccinated at the same rate at any moment in time. Rather than fitting vaccination patterns to any one country, we characterized a general pattern found in the places where polio held on the longest. That pattern involved periods of more intense vaccination that knocked down transmission followed by less intensification of efforts or even diminution of efforts after initial success. Our study uses a linear increase in vaccination rates to focus on the consequences of persisting low level transmission in the face of ongoing vaccination rather than any particular pattern in some specific location. For model settings where there is no delay in reaching the final levels of vaccination being assessed, the rate of vaccination is constant at its final level from the onset of vaccination. When there is a delay, we begin at zero and linearly ramp up vaccinations across the delay interval to a level that achieves a first infection prevalence of 300 at the end of the ramp. That vaccination level varies by the length of vaccine ramp-up, the type of waning, and the relative transmissibility of the vaccine. It is determined empirically by fitting the vaccination level to the prevalence level at the end of the vaccination ramp up. In appendix section 5, we examine how using values lower than a prevalence of 300 affects system dynamics and the inferences made in the main paper. Examples of vaccination patterns are found in figures 5 through 7.

We perform parameter sweeps over all feasible rates of effective vaccination. Our effective vaccinations thus include effects that cause live vaccines not to take as well as effects that would cause an increase in vaccine circulation as might occur with repeated emergence of cVDPVs that subsequently die out. We do not include cVDPV or IPV in our model since that would not help elucidate the mechanics of the phenomena on which we focus.

A pattern of constant levels over a long time with a subsequent ramp up was also explored. This leads to the same general conclusions we present here but it does not facilitate the identifiability inference regarding patterns of paralytic cases predicting silent circulation duration. Moreover, a ramp-up at the end generates anomalous outcomes that make comparisons across different settings difficult. For example, the prevalence may go below one before the cumulative incidence over the past year goes below the infection paralysis ratio (IPR). Patterns of vaccination vary greatly in the real world and extensive modeling work has demonstrated clearly how important capturing the past pattern of vaccinations accurately is to making valid predictions of current and future risks that arise from inadequate vaccination programs in the past (Duintjer Tebbens et al., 2014; Duintjer Tebbens et al., 2013b; Duintjer Tebbens et al., 2013c; Duintjer Tebbens et al., 2013d; Duintjer Tebbens et al., 2015; Duintjer Tebbens and Thompson, 2014, 2015; Thompson, 2013; Thompson and Duintjer Tebbens, 2014a, b; Thompson et al., 2015a, b; Thompson et al., 2013a; Thompson et al., 2013b; Thompson et al., 2013c; Thompson and Tebbens, 2012). But we want to show why these models will not adequately capture the risk of prolonged silent circulation. So we use a generic vaccination pattern that can do that.

#### Waning model rationale and description

Given the complexity of immune control of infection, waning immunity potentially has many dimensions. We choose to reduce this complexity to a single flow from the recovered (R) stage to the partially susceptible stage (P). Two parameters determine the waning process: the waning rate, and the waning depth or the fraction of transmission potential (basic reproduction number) that a population of all individuals in the P state would have compared to a population of individuals all in S. The waning depth has three components that we set to equal values. These are the relative susceptibility of individuals in the P compartment with respect to those in the S compartment, the relative contagiousness of reinfections with respect to first infections, and the relative duration of reinfections with respect to first infections. Because transmissibility is the product of contagiousness and duration, there is less waning of transmissibility than of susceptibility. The waning depth is the triple product of the fraction of waning in susceptibility, the fraction of waning of contagiousness, and the fraction of waning of duration. Since duration and contagiousness both affect transmission from reinfections, the transmissibility from the partially immune state (P) is in effect multiplicatively lowered twice while susceptibility to getting infected while in P is lowered only once.

To demonstrate the effect of different waning patterns, we chose three illustrative values of the waning rate. For each set of non-waning parameters, we then found three corresponding values for the waning depth which would result in all three parameter sets generating, in the absence of vaccination, exactly the same exponential distribution of ages of infection. We used waning parameter pair values that give a slow deep waning pattern, a fast shallow waning pattern and an intermediate waning pattern as illustrated in Figure 2 and specified in Table 1. Note that the quantity shown in Figure 2 is the product of four elements 1) the fraction of the population that would be in P state if everyone started in the R state and never experienced any subsequent infection, 2) the susceptibility of someone in the P state divided by that of someone in the S state, 3) the contagiousness of someone in the P state who gets infected divided by that of someone who got infected from the S state, and 4) the duration of infection for infections from the P state divided by duration of infections from the S state. All four of these elements determine effective reproduction numbers as shown in the appendix. Note that the differences in susceptibility between individuals in the P state and the S state does not affect the force of infection that arises from contacts of exposed individuals and determines the age of individuals experiencing their first infection. In appendix section 3 we analyze at equilibrium how model parameters affect the ratio between transmissions from first infections and reinfections.

**Figure 2:**
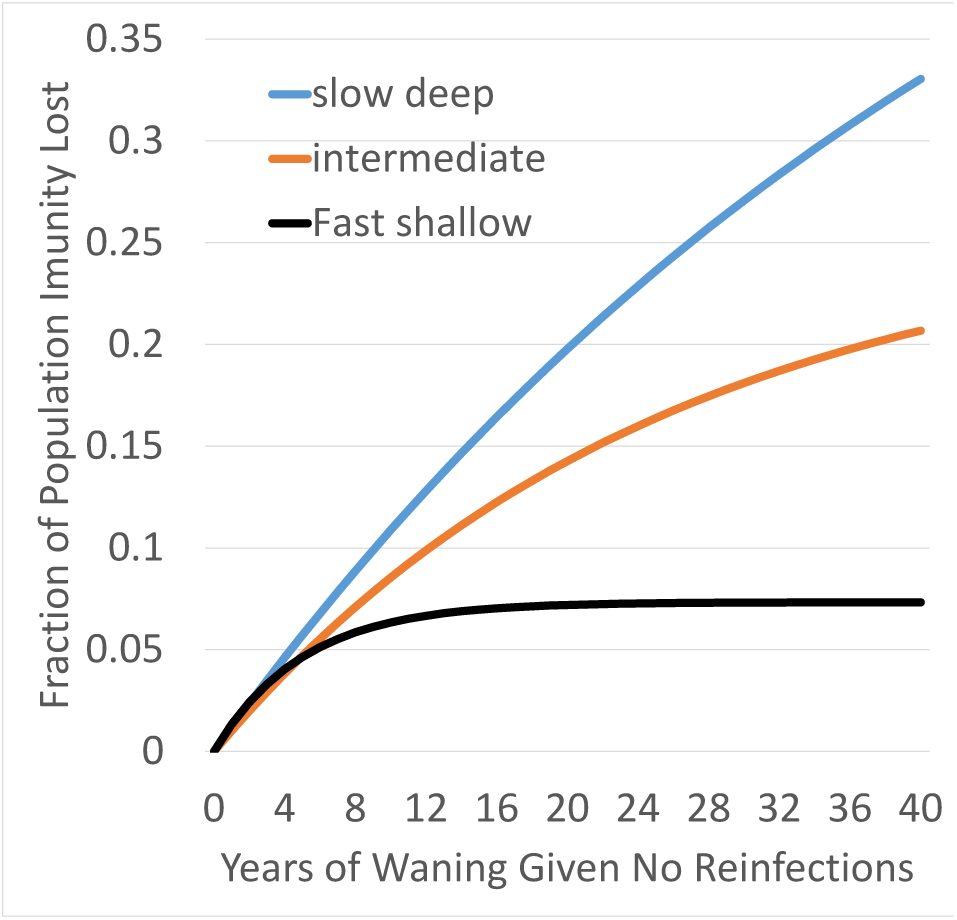
Population extent of waning for the different waning scenarios as defined by waning depth affecting susceptibility times waning depth affecting duration, times waning depth affecting contagiousness under the assumption that all three effects have the same waning depth parameters. Waning rate and depth parameters used are from Table 1.

The average age of infection in our model is determined jointly by the basic reproduction number and by waning parameters since the first generates a force of infection from first infections and the later generates a force of infection from reinfections. We examined two different average ages of infection at equilibrium before vaccination was begun: Five years for lower transmission and 3.333 years for higher transmission. The age distribution at time of first infection for all of our high transmission settings result in an *R*_0_ estimate of 15 using the methods of (Fine and Carneiro, 1999); our low transmission settings would likewise all result in an *R*_0_ estimate of 10. These two values represent high and low values for “poor developing countries” in (Fine and Carneiro, 1999) Figure 1. The basic *R*_0_ in Table 1 is the number of infections that a single infected individual would generate if all other individuals in the population were in the completely susceptible state. How we set parameter values is in the legend of Table 1. That the values in Table 1 generate dramatically less waning than the model by the Institute for Disease Modeling (Behrend et al., 2014; Wagner et al., 2014) is illustrated in Figure 7 of the fourth Appendix section.

Our primary outcome variables of interest are duration of silent circulation, defined as transmission in the absence of observed polio cases, and outcome of silent circulation. Silent circulation duration is observed in model output from the time that the cumulative incidence of first infections over the past year falls below the first Infection to paralysis ratio (IPR) until either the cumulative incidence of first infection rises once again above the IPR or the prevalence falls below an average of one individual in the population. Stable eradication at a given level of vaccination occurs when the equilibrium effective reproduction number from both first infections and reinfections is below one. The reproduction number may continue to rise for many years after the last polio case when there is slow or intermediate waning. Stability, therefore, is determined by running models for a long time. Given stable eradication, reintroduction of infection from the outside after local transmission has ceased will not result in an epidemic. With unstable eradication, however, individuals with waned immunity in the P state from figure 1 will eventually put the effective reproduction number above one.

**Table 1:**
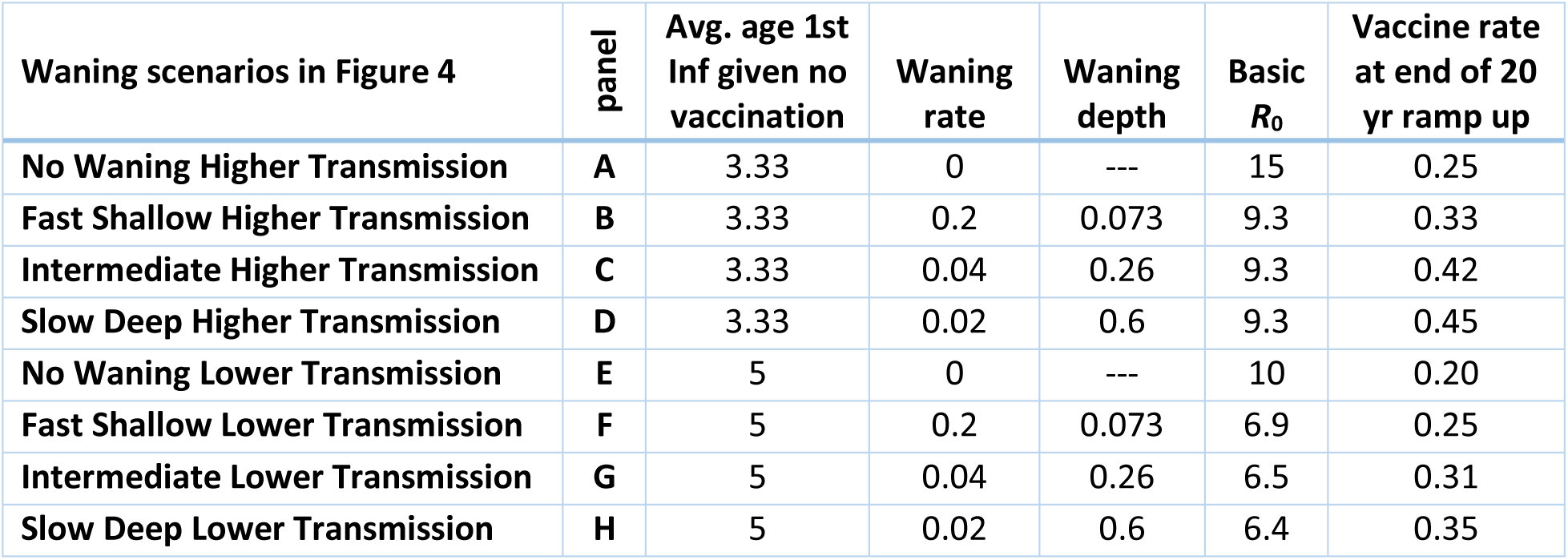
Transmission, vaccination, and waning parameters. The first three columns of numbers are set parameter values with the caveat that the waning depths for B-D were selected to give approximately the same *R*_0_ values. Then those three values were used for waning depths F-H. The basic reproduction number is set to the value found to give the desired average age at first infection at equilibrium before vaccination is begun. The vaccination rates are those that give a prevalence of infection of 300 per million at the end of the ramp up period and are found separately for each of the eight settings given the values in the first four columns.

| Waning scenarios in Figure 4 | panel | Avg. age 1st Inf given no vaccination | Waning rate | Waning depth | Basic $R_0$ | Vaccine rate at end of 20 yr ramp up |
| --- | --- | --- | --- | --- | --- | --- |
| No Waning Higher Transmission | A | 3.33 | 0 | --- | 15 | 0.25 |
| Fast Shallow Higher Transmission | B | 3.33 | 0.2 | 0.073 | 9.3 | 0.33 |
| Intermediate Higher Transmission | C | 3.33 | 0.04 | 0.26 | 9.3 | 0.42 |
| Slow Deep Higher Transmission | D | 3.33 | 0.02 | 0.6 | 9.3 | 0.45 |
| No Waning Lower Transmission | E | 5 | 0 | --- | 10 | 0.20 |
| Fast Shallow Lower Transmission | F | 5 | 0.2 | 0.073 | 6.9 | 0.25 |
| Intermediate Lower Transmission | G | 5 | 0.04 | 0.26 | 6.5 | 0.31 |
| Slow Deep Lower Transmission | H | 5 | 0.02 | 0.6 | 6.4 | 0.35 |

### Model equations and numerical solutions

The model equations are presented in the appendix along with the code used to implement those equations in Berkeley Madonna (Oster and Macey, 2015).

### Model analysis

The model is analyzed both mathematically and numerically. Numerically the model is run to WPV equilibrium before introducing OPV vaccination. Numerical analyses were checked to see that they were not affected by the numerical integration algorithms used or the time steps employed. Mathematical analysis is presented in Sections 2-4 of the appendix.

## Results

### Model generated patterns of drop in first infection prevalence

Figure 3 shows the drop in incidence of first infections during a 20 year ramp up delay of vaccination for parameters in Table 1. Given the infection to paralysis ratio of 1000 used in this analysis and the incidence of first infections of about 3,860 per year at the end of the ramp up, 3-4 polio cases per year in our population of one million would be expected at the end of the ramp up. The consequences of using other prevalence values at the end of the vaccination ramp up period besides the 300 value used here are presented in appendix section 5.

The procedure for setting parameters outlined in the Table 1 legend was chosen to make the force of infection identical at both the endemic equilibrium and the point at the end of the ramp up, for all waning scenarios, at a given average age of first infection. By doing this, we demonstrate that the pattern of polio cases over time in a population cannot be used to determine the waning pattern in that population. That inference is important because in the next section we will show that waning patterns with identical effects on the population level of infection before vaccination is begun have huge effects on silent circulation duration after the last polio case. Adding information about effective vaccination levels might theoretically help determine waning patterns. From table 1 we see there is some increase in vaccination level needed to get to our fixed prevalence of first infections as we progress through fast-shallow to slow deep waning scenarios. But given the difficulties in determining what the effective levels of vaccination have been in populations over time, the waning patterns will not be identifiable from standard polio program data.

**Figure 3:**
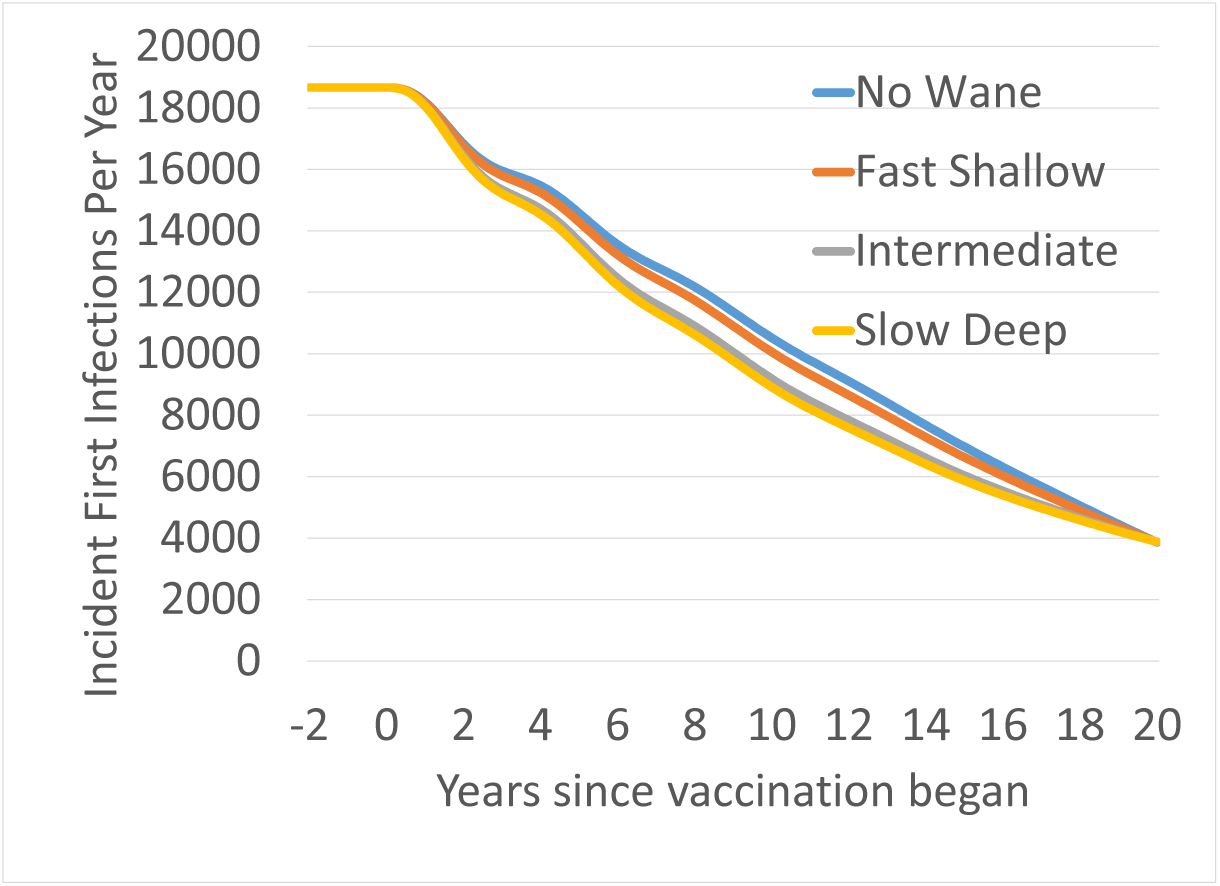
First infection incidence as a function of waning scenario across a 20 year delay period during which first infection prevalence is reduced to 300 per million for the higher level of transmission. Settings used are those for a type 3 virus and settings A-D from Table 1.

### Silent circulation patterns

Silent circulation begins in our model when vaccination brings down cumulative incidence of first infections over the past year to less than the first infection to paralysis ratio (IPR). That point is a likely time when the last polio case might be detected. Silent circulation ends with either stable eradication, unstable eradication, or recurrence of polio cases. We declare eradication when the prevalence of infection goes below one. If the cumulative incidence of first infections since the last polio case rises above the IPR before the prevalence goes below one, then we declare a recurrence of infection.

Figure 4 presents silent circulation duration by levels of vaccination and indicates by color whether the silent circulation ended with a recurrent polio case, unstable eradication where a new introduction could lead to a new outbreak, or stable eradication where outbreaks would be small. The level of vaccination in Figure 4 is the rate of vaccine takes and for completely susceptible children this is assumed to be the same across the age range getting vaccinated. In the first curve in each panel there is no delay in vaccination reaching its final levels. In the second there is a 20 year delay during which vaccination is ramped up as per Table 1 generating drops in paralytic cases following Figure 3 patterns. The reader will better understand Figure 4, when they understand how the dynamics in figures 5 and 6 generate Figure 4 patterns.

**Figure 4:**
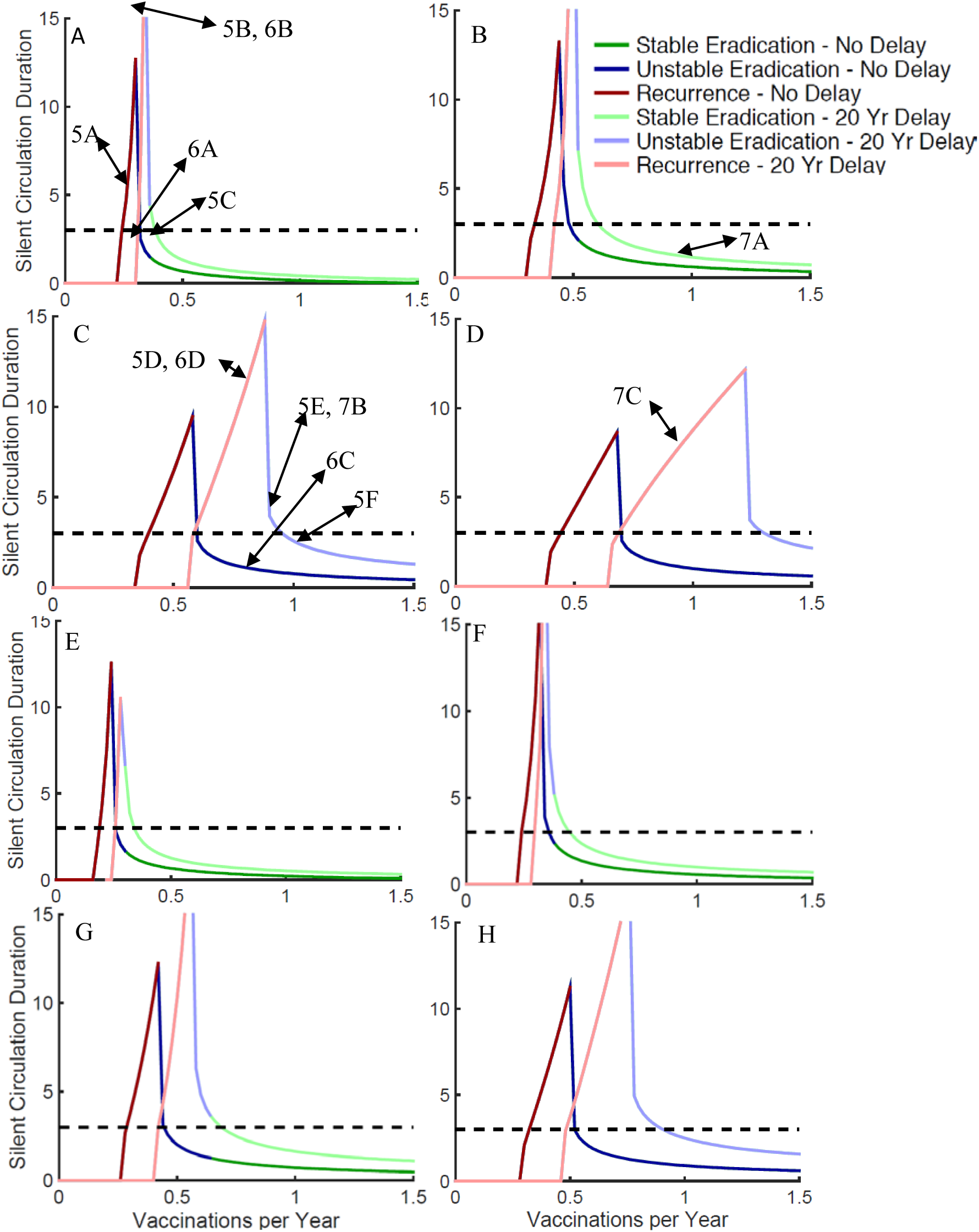
Patterns of duration of silent circulation as a function of final total vaccination rates in less than five year olds by waning immunity scenarios: A&E) no waning, B&F) fast shallow waning; C&G) intermediate waning; and D&H) slow deep waning. A-D have an average age of first infection of 3.33 years while E-H have an average age of 5 Type of silent circulation (stable, unstable, and recurrent) is indicated by color. Horizontal dashed line illustrates 3 year silent circulation. Parameter values used are shown in Table 1. The points on these curves where the dynamics are illustrated in subsequent Figures are indicated by Figure number and Panel letter.

Figure 4 reveals 3 effects as waning progresses from no waning to slow deep waning across panels A->D and E>H. As waning becomes slower and deeper while keeping the effects of waning at equilibrium in the absence of vaccination, 1) it takes higher final vaccination rates for eradication to be achieved. 2) The difference in vaccination rates required for eradication instantly achieving the final vaccination levels and a 20 year delay increases. 3) The breadth of vaccination levels that result in prolonged silent circulation increases.

All of these differences are greater when transmissibility is at the upper end of the plausible range for higher transmission countries (A->D) than when it is at the lower end of that range (E->H). The differences between no waning and fast shallow waning are small. In these scenarios there is no (meaningful) ongoing waning after 5 years, so that there is no ongoing accumulation of people susceptible to reinfection after 5 years. As seen in Figure 2, there is such an ongoing accumulation of partially susceptibles with intermediate waning, and there is twice as much accumulation with slow-deep waning.

A big determinant of how likely we are to encounter prolonged silent circulation in the real world is the width of the vaccination levels across which there is prolonged silent circulation. If vaccination levels have to be held in a narrow range in order for there to be prolonged silent circulation, then natural fluctuations will eliminate that risk. Thus one conclusion from figure 4 is that prolonged silent circulation will become increasingly likely in the real world as the delay in achieving effective vaccination levels increases and as the amount of ongoing waning increases. These two factors are more important than the level of transmissibility in determining silent circulation risks. The effects of these factors, however, are greater at higher levels of transmissibility.

This is important because empirical experience with prolonged silent circulation risks is based on experience in settings with lower transmission levels and shorter times between beginning vaccination efforts and ending polio cases than are characteristic of the currently or recently endemic countries. Thus there is reason to expect that empirical experience may not be a good guide for setting the time period after the last polio case at which we are confident that there is no ongoing silent circulation.

A final thing to note in Figure 4 is the effect of waning scenarios, transmission levels, and ramp-up times on whether eradication is unstable or stable. Stable eradication in Figure 4 occurs with no waning (Panels A&E) or fast shallow waning (Panels B&F). For waning that continues beyond 5 years, however, it only occurs at the lower bound of high transmissibility conditions and the lower depth of waning (panel G).

### Effective Reproduction Number analyses to explain silent circulation patterns

The total effective reproduction number for our model is the sum of the effective reproduction number for first infections plus the effective reproduction number for reinfections in each of the age groups. The mathematical logic supporting this is shown in appendix section 2.

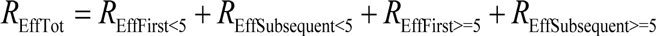

The dynamic pattern of these four components helps determine whether a given vaccination level will result in stable eradication, unstable eradication, or a recurrent case of polio. The total time and extent to which the effective reproduction number stays below one determines how fast and how far the prevalence of infection will fall. The effective reproduction number when vaccination is first begun always starts out at the value of one in all of the dynamics panels in Figures 5 through 7 because we start vaccination only when the system is at equilibrium.

Note that in Figure 5 before and during the 20 year ramp up, all panels are identical within the two waning scenarios examined (A-C no waning and D-F intermediate waning). During the 20 years of vaccination ramp up, the total effective reproduction number dips only slightly below one. For the no waning scenario in panels A-C of Figure 5 there are only first infection events so the above equation has only two elements on the right hand side. During the ramp up delay, vaccination reduces the contribution of first infections in the under-five age group. But that drop is compensated for by more children being susceptible at their fifth birthday so that the greater than five-year-old susceptible group contributes more to the effective reproduction number.

Panels D-F show a similar pattern of the total effective reproduction number staying near one during the ramp up for intermediate waning. But the compensation behind this differs. The decrease in the green area of first infections in children under age 5 is mostly compensated for by an increase in transmission from reinfections in the older age group that is not receiving vaccinations. There are fewer individuals reaching age 5 still susceptible (purple color) because there are fewer fully susceptible children under age 5 to reach that birthday and because reinfections continue to increase as polio levels fall so there is more force of infection overall to infect the over five age group.

Another difference between the no waning and intermediate waning scenarios is the size of the vaccination boosts after the delay that it takes to get eradication to occur within the three year period used to declare eradication. In the absence of waning, it takes very little increase in the size of the boost to go from just enough vaccination to eliminate polio cases for a year to enough vaccination to stably eradicate transmission in less than three years. Given intermediate waning, it takes a big boost. That is because the boost is mainly affecting the under-five age group that is directly vaccinated. We will see later in Figures 7 and 9 the importance of effects of indirect vaccination through transmission of the vaccine. These effects are of course acting in Figures 5 and 6 as well. But here the reader should note the great rise in the yellow area of reinfection effective reproduction number that compensates for the fall in the reproduction number from the green age group receiving vaccination when a vaccination boost is experienced. Also to be noted is that the rise in the reinfection reproduction number continues even after eradication such that eradication quickly becomes unstable as the total effective reproduction number goes above one.

In Figure 6 we see how a delay affects the dynamics in the absence or presence of waning. Both in the presence and in the absence of waning, a delay causes a decrease in the size of the drop in the total effective reproduction number when vaccination is boosted. In both cases that decrease occurs because although the contribution of the group being vaccinated has been diminished during the delay, increases in the contributions from other groups have largely compensated for this decrease. When there is no waning, it is the contribution from the older age group who did not get vaccinated that increases during the delay. With ongoing waning, it is primarily the contribution from older individuals whose immunity has waned that increases. The other major difference is that when there is ongoing waning, it takes much higher levels of vaccination to get elimination of polio cases and eradication of transmission.

Note that the difference in the compensating groups implies different policies for addressing a failure to eliminate polio cases or a recurrence of polio cases after their elimination. If there is no waning, then the increased susceptible individuals over age 5 will all have accumulated during the delay and thus young people should be targeted for special control programs. But in the presence of ongoing waning, it is the age groups that have passed the most time since vaccination or natural infection whose immunity is most likely to have waned. Thus older, rather than younger, age groups should be targeted.

**Figure 5:**
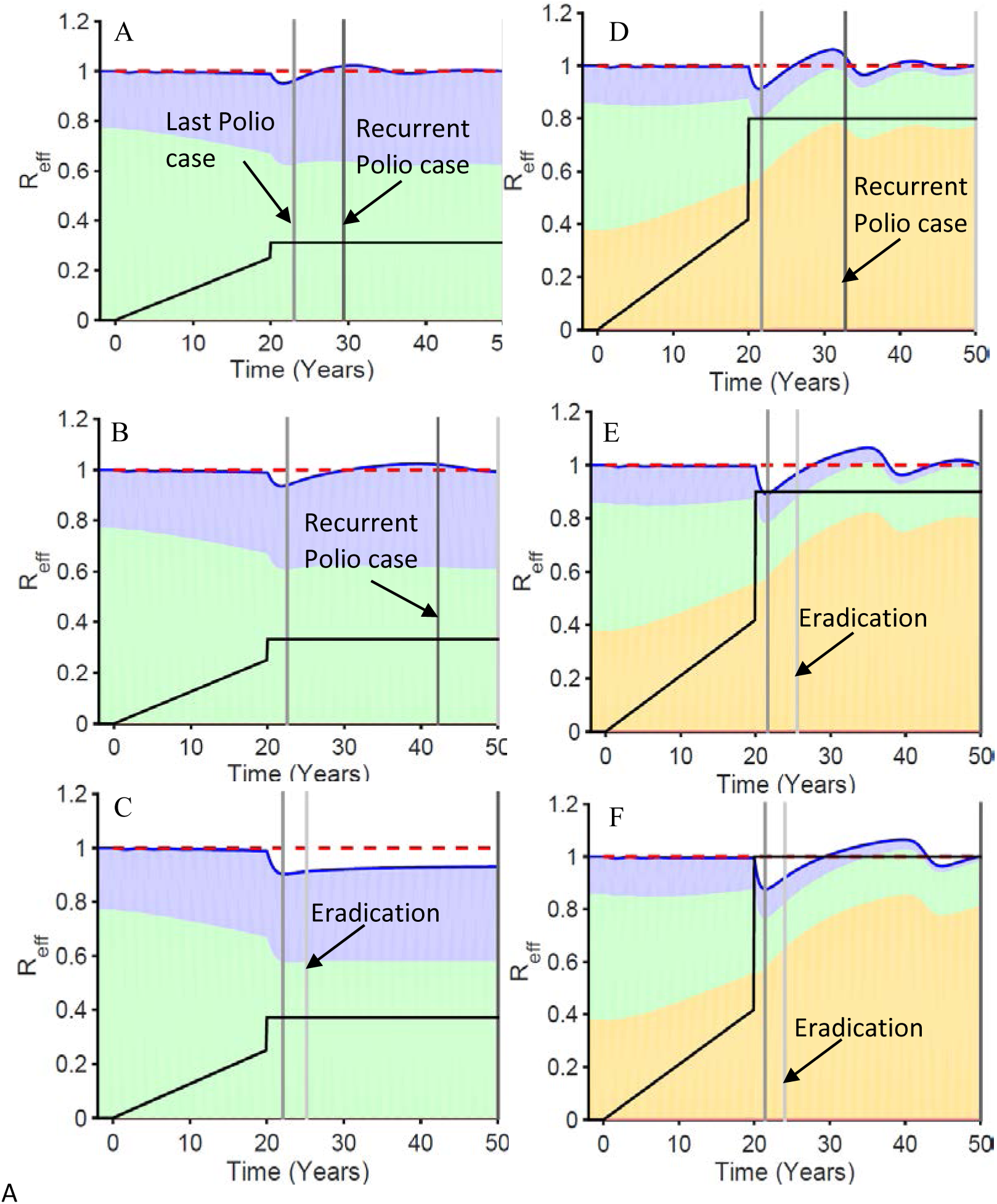
The dynamics of the effective reproduction number and its components as vaccination levels are increased. Intermediate waning, the higher level of transmission, a 20 year delay and serotype 3 settings were used. The position of each panel on Figure 4 curves are indicated on Figure 4. The final vaccination levels for panels are A=0.32 effective vaccinations per year, B=0.34, C=0.38, D=0.8, E=0.9, F=1.0. The black vaccination level lines ramping up to 20 years and then jumping to the final vaccination level use the same left hand scale as the effective reproduction numbers. The blue shaded area is the contribution to the effective reproduction number from first infections in the five and older age group, the green is first infections from the under 5, the yellow is reinfections in the five and older, and the red (barely visible below the yellow) is from reinfections in under age five. The dark blue line summing all shaded areas is the effective reproduction number. The red dotted line is the transmission threshold. The time of eradication, last polio case, and a recurrent polio case have increasingly darker grey shading.

**Figure 6:**
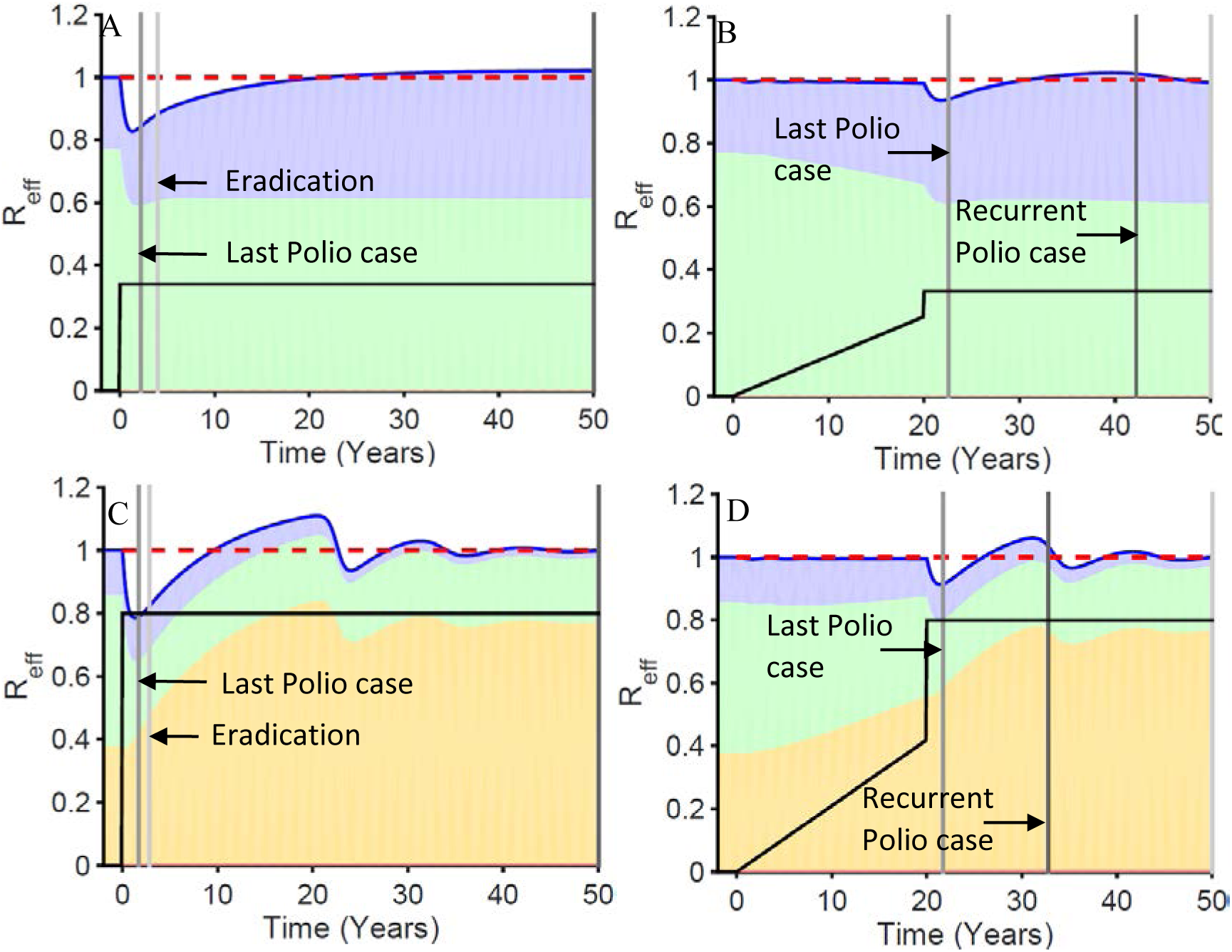
The dynamics of the effective reproduction number and its components with and without a delay in reaching final vaccination levels. Intermediate waning, the higher level of transmission, and serotype 3 settings were used. The positions of each panel on silent circulation duration curves are indicated on Figure 4. The shading and lines have the same meaning as Figure 5.

Figure 7 shows how waning patterns affect the outcome of eradication efforts given nearly identical patterns of decrease in polio cases as shown in Figure 3. All three panels have waning that produces the same average age of first infection before vaccination is begun given the same level of transmissibility as shown in Table 1. We focus on three times on the horizontal axis of each panel. In the time before vaccination is begun, all three waning scenarios have the same fractions of age and immunity groups that contribute to the total endemic effective reproduction number of one. During the ramp up, in order to get the nearly identical drop in first infections, however, it takes a higher level of vaccination at the end of the ramp up in those scenarios with higher levels of ongoing waning many years after infection. That means that at the time of the vaccination boost, as we go from fast shallow to slow deep, the contribution of reinfections to the effective reproduction number increases and contributions of the vaccination target group decreases. Finally we look at the response to the boost given the same level of vaccination. The difference between those scenarios with and without ongoing waning is dramatic. There is very little rise in the contribution of reinfections for the fast shallow scenario and consequently the effective reproduction number drops dramatically and eradication is achieved rapidly and stably. In contrast, the rise in the contribution of reinfections is much higher in the intermediate and slow deep scenarios, and in both of these, eradication is achieved slowly if at all, and unstably. The difference between intermediate waning and slow deep waning is much less. But the greater increase in the contribution of reinfections to the effective reproduction number in the slow deep scenario compared to the intermediate scenario makes the difference between unstable eradication and recurrence.

**Figure 7:**
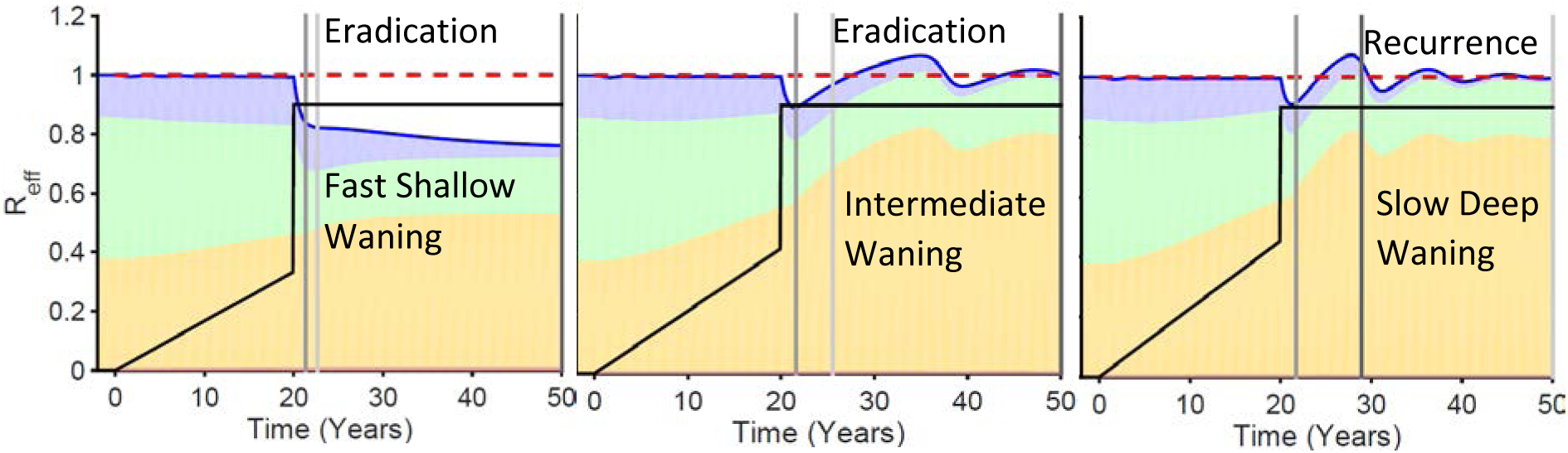
Changing dynamics of the elements of the total effective reproduction number across different waning scenarios explain the effects of those waning scenarios when final total vaccination levels are kept constant. Higher transmission parameters along with IPR and OPV/WPV transmissibility characteristic of type 3 were used.

**Figure 8:**
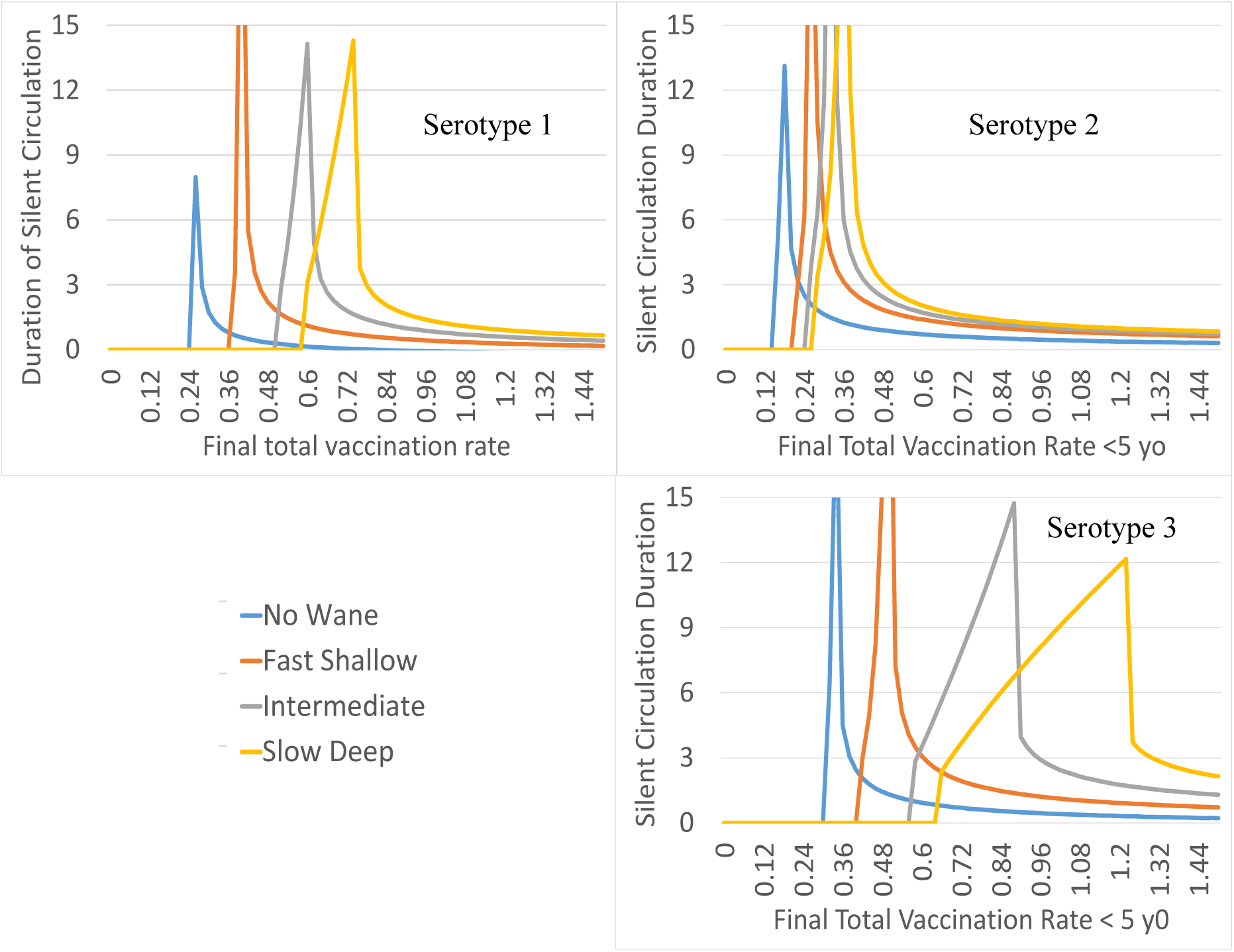
Length of silent circulation by final total vaccination rates in less than five year olds and waning scenarios for different serotypes. Settings include a 20 year ramp up time delay and high transmission.

### Serotype differences in silent circulation potential

In this analysis we assume serotypes differ only in their first infections to paralytic infections ratio (IPR) and in their OPV transmissibility to WPV transmissibility ratio (OWR). We use values of OWR of 0.37, 0.57, and 0.25 and values of IPR of 200, 2000, and 1000 for types 1, 2, and 3 respectively. These values are in the range used by other modelers (Duintjer Tebbens et al., 2013c). Based on our simulations, Serotype 3 is the hardest to eradicate and Serotype 2 the easiest (Figure 8). The risks are smaller for the no waning scenario and the fast shallow waning scenario and the changes in silent circulation for these first two waning scenarios are mostly in the level of vaccination where prolonged silent circulation is seen rather than the width of the vaccination range across which it occurs. But for intermediate and slow deep waning the effects on the width of the vaccination interval resulting in prolonged silent circulation is quite large.

**Figure 9:**
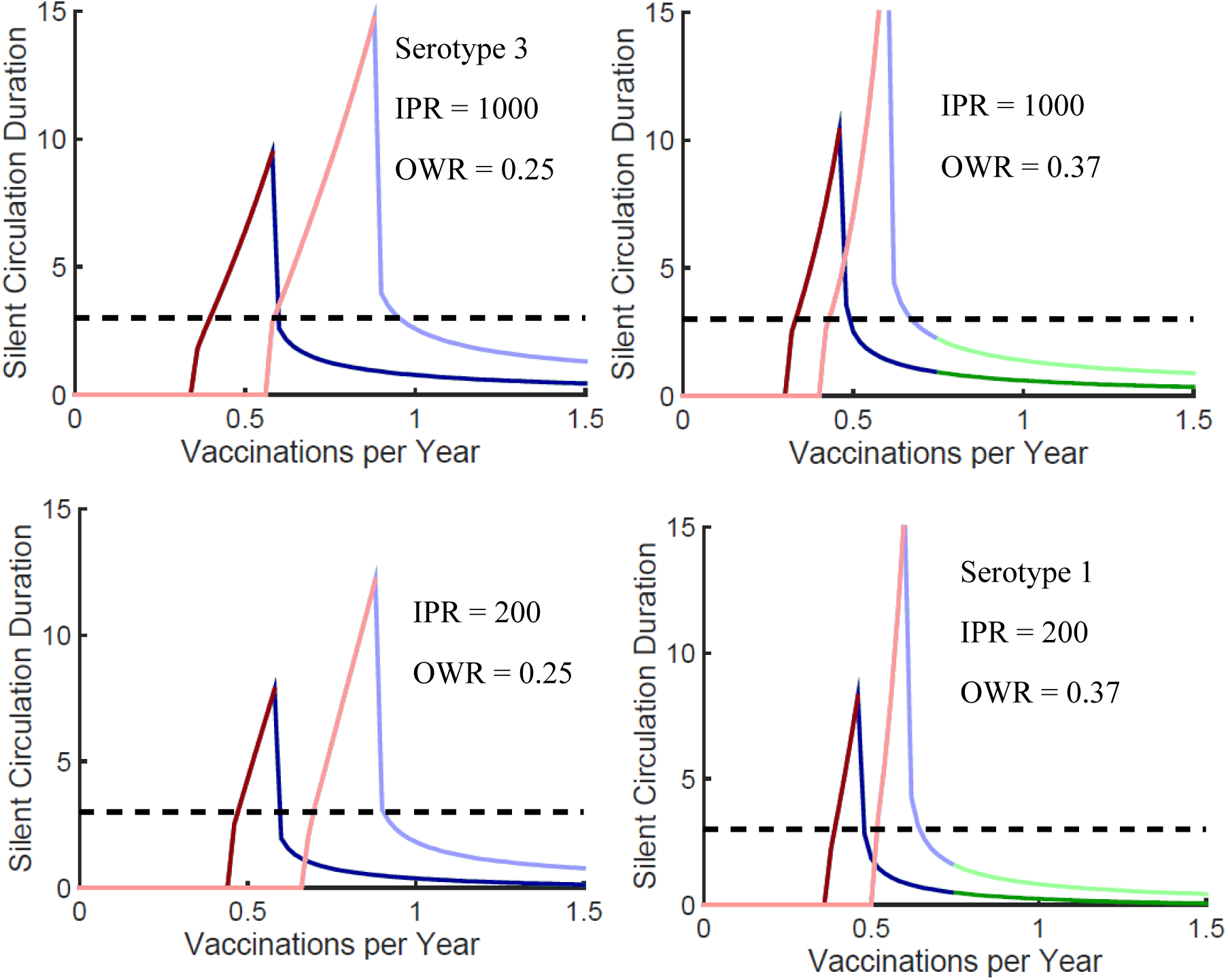
Comparison of the effects of changing just the infection to paralysis ratio (IPR) or just OPV/WPV transmissibility ratio (OWR) to make the transition from serotype 3 settings to serotype 1 settings for intermediate waning settings with no delay and twenty year delay before instituting final total vaccination rates per year in less than five year olds.

To separate the effects of the IPR and the OWR we compared settings starting with the serotype 3 settings of 1000 and 0.25 and changed them to 1000 and 0.37 and 200 and 0.25 so that we could see more clearly what leads to the difference between serotype 1 and serotype 3. In Figure 9 we see that the big effect on the risk of silent circulation is the lower vaccine transmissibility rather than the higher IPR. The lower vaccine transmissibility causes a sharper rise in the contribution of reinfections to the total effective reproduction number when vaccination is boosted to its final level. Changing the IPR does not change the dynamics of how the different elements contribute to the overall effective reproduction number. It just changes the time of the last polio case and the time of recurrence.

### Levels of prolonged silent circulation

Prolonged silent circulation that eventually results in eradication in the situations we examined had a maximum prevalence of wild polio virus of 110 per million or lower for the slow deep waning and 81 for the intermediate waning. Those levels were highest at the time of the last polio case and decreased progressively with time.

For the scenarios that ended in a recurrent case, the maximum prevalence occurred when the recurrent case appeared right after three years. For both slow deep and intermediate waning the prevalence in this case was 200 per million at the time of the last polio case. It went down for a year, and then came back just before three years to 500.

These levels would take extensive environmental surveillance to detect. Not only would environmental surveillance need to be able to detect a single reinfection in every population of 10,000, but the detection sensitivity would have to be more than 10 times greater than that needed to detect a first infection. Thus environmental surveillance as currently being implemented is likely to miss the prolonged low level silent circulation that we have illustrated.

### Levels of vaccination in older age groups needed to avoid prolonged silent circulation

To evaluate the effect of vaccinating older age groups before OPV use is stopped, we determined the rate at which the five and older age group would have to be vaccinated at the time of the boost in vaccination levels in order to bring about eradication in less than three years from the last polio case. The worst case situation is the high transmission scenario with slow deep waning after a 20 year delay that only reaches a first infection prevalence of 300 per million and where the final vaccination level after the boost is just enough to generate a three year delay before a recurrent case appears. In this situation only 8% of the ≥ 5 population needs to be vaccinated per year in order to get eradication within three years of the last polio case. At higher final vaccination levels of the < 5 population it takes less. If the ramp up goes to a lower prevalence of polio before the vaccination is boosted to final levels, it also takes less vaccination in the older age group to insure that silent circulation is not prolonged.

Since the model analyzed here does not break down the ≥ 5 population into adults and older children, we cannot say what age groups are the most important to vaccinate in order to boost waned infections. But that would seem to be adults and not older children since the adults will have experienced more time since getting directly vaccinated with OPV and have less exposure to children who have been recently vaccinated.

But when vaccination of the older age groups is only during the third year after the last polio case, it takes considerably more vaccination of the older age groups to eliminate prolonged silent circulation. In our worst case scenario discussed above, it would take nearly 1.4 vaccinations per adult per year. But if vaccination of the older age groups occurred during the second and third years after the last polio case, only 29% of the older age group would need to be vaccinated per year.

## Discussion

### The major message in our findings

Our theoretical model analysis demonstrates that small amounts of waning immunity could lead to prolonged silent circulation of WPV without paralytic polio cases. All the waning scenarios in our model cause less transmission from reinfections in the pre-vaccine era than do the models of the Kid Risk group (Duintjer Tebbens et al., 2013c). The levels at which such prolonged circulation occurs in our models are low and might be hard to detect with environmental surveillance. If waning stops after five years, as in the models of the Kid Risk group (Duintjer Tebbens et al., 2013c), the risks of prolonged silent circulation are small. Ongoing waning after five years together with long periods during which WPV circulation has been suppressed but not eradicated increase the range of past vaccination histories that can generate prolonged silent circulation. This increases the chances of prolonged silent circulation. High transmission potential makes the emergence of prolonged silent circulation more likely as it reduces the amount of waning needed. In our models, notable risks of prolonged silent circulation or unstable eradication do not emerge across the parameter ranges we examined at the level of transmission estimated by [Fine and Carneiro] for “good industrialized countries” before 1950. Those levels of transmission should be considerably less today. Of course pockets of poor hygiene and vaccination could still cause problems anywhere.

It is transmission from previously infected or immunized individuals that sustains silent circulation in our models. Adults who were previously infected but who have escaped contact with WPV or OPV infected children for a long time have the best chances to sustain silent circulation. We did not separately examine this adult group in the analysis presented here. Lower transmissibility of OPV viruses is consequently a major factor increasing the risk of prolonged silent circulation. That is because in current eradication scenarios, it is mainly transmission of OPV that boosts immunity in adults. Consequently, type 3 WPV carries the highest risk of prolonged silent circulation because its vaccine is the least transmissible. This prediction of which serotypes are going to have the longest silent circulation is different from the predictions made in (Kalkowska et al., 2015). That is because in the Kid Risk model used in that analysis (Duintjer Tebbens et al., 2013c), only a fast-shallow waning formulation with no ongoing waning was used. Thus the infection to paralysis ratio became the dominant determinant of silent circulation duration and type 2 had the longest silent circulation in places like India because it has the highest infection to paralysis ratio.

A key finding is that the nature of this phenomenon is such that it could be completely missed by models fitted only to polio case data. Figure 3 shows that the different waning scenarios that have such dramatic effects on the duration of silent circulation in Figure 4 have only small differences in first infections that can lead to polio cases. These differences in Figure 3 would not be detectable in reality. In other words, we show that nearly identical patterns of decreasing cases on the path to polio elimination are consistent with models having either low or high risks of prolonged silent circulation. Given expected noise from imperfect surveillance and chance events, these two findings indicate that models fit to polio case data will have little or no practical capacity to predict how long silent circulation will continue after the last polio case.

The levels of vaccination that finally achieve polio case elimination in our model analysis are lower than the levels needed for case elimination in India, Nigeria, Pakistan, or Afghanistan. That is true even when our effective vaccination rates are divided by their low vaccination take rates (Duintjer Tebbens et al., 2013a) to turn them into actual vaccination rates. Another factor that could stretch our Figure 4 curves to the right is the emergence of cVDPV. As seen in Figure 4, most prolonged low-level silent circulation in our model ends with the recurrence of a new observed case of polio. If VDPVs evolve to circulate longer than OPV does, than the resulting cVDPV will lower the chance of recurrence and extend our curves to the right. Additionally, our vaccination pattern model lowers the vaccine rates needed for eradication as the prevalence level used to set the endpoint for the ramp up is raised. The mechanism behind this is that at lower prevalence settings at the end of the ramp up, there is a lower fraction of the force of infection that derives from fully susceptible children. Thus a higher rate of vaccination is needed to lower the total force of infection since the big effects come from vaccinating fully susceptible individuals and there are fewer of them. Thus the total effective reproduction number must be reduced by more OPV being transmitted from the age group receiving vaccine to the older age groups where immunity has waned.

Models by the Kid Risk group (Duintjer Tebbens et al., 2013c) that lead to short duration of silent circulation before eradication have had a dominant influence on polio eradication policy. These models have been useful in guiding vaccination policies to achieve elimination of polio cases. The success of these models for that purpose most likely lie in the fact that polio case data has some power to identify patterns of immunity deficiencies arising from failures to effectively vaccinate all children using the Kid Risk model (Duintjer Tebbens et al., 2013c) and from the fact that failure to reach children with vaccination is strongly correlated with failures to eliminate polio cases. This success, together with a history of struggles by polio eradicators with naysayers who have brought forth many objections to pursuing eradication because of theoretical difficulties, has inured the leaders of polio eradication efforts to warnings like those raised by our work.

But our work does not make us naysayers of eradication. Polio is the greatest global communitarian effort ever undertaken by mankind. Its success is crucial for advancing further such efforts. We only bring a message that action should be taken to insure against silent circulation when OPV use is stopped. In our model, only modest levels of vaccination in adults are needed for such insurance. Adult vaccination is needed only for relatively brief periods before OPV cessation. Vaccination of adults before OPV cessation would also reduce the risk of circulating vaccine derived polioviruses (cVDPV) emergence during cessation. cVDPV emergence risk has been a major focus of endgame modeling (Duintjer Tebbens and Thompson, 2015; Famulare et al., 2015). It is likely to occur more frequently after OPV cessation than the prolonged low-level silent circulation on which this paper focuses. But a WPV recurrence due to silent circulation at the time of OPV cessation will require more extensive control efforts and have greater risk of getting out of control due to the higher transmissibility of WPV and due to the role of adults who are no longer getting their immunity boosted by OPV transmission from vaccinated children.

### Other modeling studies showing low risk of prolonged silent circulation

Two studies have concluded that the risks of prolonged silent circulation are fairly circumscribed (Famulare, 2015; Kalkowska et al., 2015). But they have not fully addressed the dynamics that we have illuminated with respect to the range of waning possibilities and they fitted their models to polio case data which we show here to be uninformative about silent circulation after the last polio case as illustrated by Figures 3 and 4.

(Famulare, 2015) assumes that dynamics affecting silent circulation stay constant in the years before the last polio case because the effective reproduction number stays relatively constant. But Figures 5-7 show that the shifting composition of susceptible individuals that stabilizes the overall effective reproduction number is the major determinant of whether or not there will be prolonged silent circulation after the three year period used to confirm eradication.

(Kalkowska et al., 2015) builds on the basic Kid Risk model (Duintjer Tebbens et al., 2013c) and thereby specifies waning dynamics that are even faster although somewhat deeper than those in our fast shallow waning scenario. The model used in this study has been found consistent with case occurrence and vaccination data in a variety of circumstances (Duintjer Tebbens et al., 2013c). But there are various problems in accepting the conclusions of this study. One is that we show that fitting case data and vaccination data is not enough to predict the risk of silent circulation, as a wide range of waning settings yield very similar case by vaccine level trajectories. Another is that the waning pattern used in this analysis is inconsistent with expert opinion (Duintjer Tebbens et al., 2013a, b). Long term waning patterns, however, are poorly studied, so expert opinion may not count for much. Antibodies against other infections wane in phases with progressively slower but persistent waning (Teunis et al., 2016). Given that polio reinfections are not a threat to health, there is little reason for the immune response to polio to have evolved such that there is no ongoing waning.

A further issue is that the model used by (Kalkowska et al., 2015) has many parameters that are adjusted to fit observed infection patterns and the major reason very fast waning with no ongoing waning was adopted was because fast shallow waning best fit infection patterns (Duintjer Tebbens et al., 2013c). The model complexity precludes a thorough analysis of dynamic effects of model components. Thus it is possible that unfortunate rigidities were built into the model that inappropriately precluded model fits with ongoing waning. Additionally, the total amount of waning in this model is larger than in our waning models in terms of the effects that waning would have on the average age of first infection at equilibrium in the absence of vaccination. That could be causing all the recent Kid Risk models, including (Kalkowska et al., 2015), to lower transmission parameters below levels that actually occur. Similarly, it could be causing them to be only able to fit observed polio case data by having waning completely stop after five years. The total amount of early waning was set by expert opinion. Thus it seems possible that because early waning was set too high, ongoing waning had to be inappropriately eliminated. It would help to resolve these issues if the exact data fit and the computer implementation of the model that was fit were published so that possible fits could be explored by a wide community of scientists. This level of openness, which the scientific community is moving towards (Nosek et al., 2015), would help a growing community of modelers using diverse and innovative methods to contribute to important decisions about OPV cessation.

### Studies needed to determine waning immunity effects

Widespread hidden persistence of low levels of silent WPV transmission when OPV is stopped would be a devastating problem. It would more urgently require reversion to OPV use than would cVDPV emergence because WPV are more transmissible than cVDPV, at least in the models that have addressed the emergence of cVDPV (Duintjer Tebbens et al., 2013c). An assessment that this risk is low and that changing the message to the public that only children need to be vaccinated would be harmful to the program makes the Global Polio Eradication Initiative resistant to the idea of vaccinating adults. It is this resistance that makes further work on determining the effects of waning on transmission so urgent. This resistance will only be overcome by solid scientific evidence that waning immunity could generate the risks of prolonged silent circulation that arise in our model.

There might be a chance that all waning ceases after five years without any boosting by exposure to live polio viruses. Perhaps repeated boosting will have made older people resistant to any waning of immunity once such repeated infections stop. But the basis for this hope is weak. Immunity waning against non-paralytic polio infection transmission has been mainly addressed by administering OPV and observing excretion levels and duration. The demonstration of complete loss of immunity to this high dose administration in elderly adults with evidence that their immunity had been primed years before is concerning (Abbink et al., 2005). Also concerning are the demonstration of waning between ages 5 and 10 (Jafari et al., 2014) and the waning of immunity to vaccine doses modeled by (Behrend et al., 2014; Wagner et al., 2014). But these demonstrations of waning are to extremely high doses given over a very short time. A theoretical framework has been presented that explores how immunity to such short high doses can differ greatly from more realistic exposure patterns (Pujol et al., 2009). That theoretical framework might provide a basis for further explorations as to the relative rates of waning to high and low doses of exposure.

A more empirical framework for exploring the effects of exposure dose on waning is provided by a model of immune processes that integrates data on antibody waning with a beta-Poisson distribution of infection risk by dose (Behrend et al., 2014; Wagner et al., 2014). We analyze the Behrend-Wagner model for what it has to say on dose relationships for waning in appendix section 4. Overall, the Wagner analysis indicates that there is considerably more ongoing waning than there is in our model at the parameters we have presented. See Figure 7 in section 4 of the appendix. Just as with our model, to make predictions from the Wagner model about silent circulation risks will require appropriate fitting to data and there is no empirical base for inferring the effects of low dose exposure.

An even more empirically based model of antibody level waning has recently been analyzed by (Teunis et al., 2016). The analysis, however, is for pertussis data rather than polio data. It shows that for pertussis an early fast waning phase is followed by a slower waning phase. This is probably a more realistic model than our one phase model. Data for the second phase of waning, however, may be contaminated by hidden boosting. If any saved sera across age groups and time in populations from individuals who would have had original infections from OPV or WPV and thereafter only IPV were used in the population, that would at least allow for a better description of antibody waning.

Given an area of ignorance of such great potential importance to polio eradication costs and success, it seems important to use either the theoretical framework of (Pujol et al., 2009) or the more empirical approach of (Behrend et al., 2014; Wagner et al., 2014) to design studies of dose effects on waning immunity before cessation of OPV use makes such studies impossible. A a range of vaccine doses from low to high could be given to adults in populations with high transmission and long periods of suppressed transmission without eradication. The observation of responses to such administration in the manner of (Jafari et al., 2014) should help fit models that can better predict the risks of prolonged silent circulation.

Another approach to pushing back our ignorance about waning might be to use already collected data from polio eradication efforts. The sequences collected from polio cases over time might reveal something about the patterns of contributions to the force of infection by first infections and subsequent infections over time. The length of the transmission path from one polio case to another increases as more steps in that path involve reinfections. The overall pattern of all distances between all sequences as a function of time might reflect the increasing role of reinfections over time. But experience with HIV (Volz et al., 2013) suggests that in addition to sequences it might be necessary to integrate other information reflecting the force of infection. Otherwise the time scaling in fitting evolutionary processes could eliminate the relationship being sought. The joint time patterns of distances between all samples, polio cases, and the average age of polio cases could provide a signal to which transmission models could be fit to estimate waning parameters. A valuable dataset to analyze is the pattern of documented cases and sequences from India. There seems to be little reason not to make this data available to all who can analyze it productively. The technology for extracting useful parameter estimates and useful inferences from such data are evolving so rapidly and diversely that restricting that data to one group seems counter-productive.

If OPV is stopped worldwide and detectable transmission of WPV or cVDPV emerges, there could be one last chance to assess waning by thoroughly documenting immune levels and infection rates across all age groups. Such studies would be expensive because they need to detect low levels of infection across all age groups and therefor require extensive stool collections and examinations. Also detecting excretion levels after vaccination across all age groups will be expensive. But determining the risk of silent circulation being present in other populations will be an urgent priority that would justify the expense. Let us hope that we do not have to reach such urgency in such a difficult situation in order to decide to illuminate the effects of waning immunity on the eradication endgame.

The time for performing studies that require assessing immunity by administering OPV is quickly running out because OPV use is being stopped. OPV2 use has already been stopped globally except in Borno state in Nigeria where a cVDPV2 was found in sewage about the time that vaccination with type 2 was stopped. OPV1 and OPV3 will be stopped globally three years after the last detection of either in polio cases. Therefore, other strategies for assessing and controlling the risks of prolonged low-level silent circulation are needed.

Environmental surveillance (ES) offers a strategy for assessing silent circulation risks that can be used both before and after OPV cessation. The current state of environmental surveillance the plans for expanding it are described in (GPEI, 2015). The observations made in this paper have implications for how ES could be most informatively intensified and how its results most informatively interpreted. The current use of ES data is just to indicate whether silent circulation exists. ES could also be used to indicate what model scenarios that generate different patterns of silent circulation rates are the most likely. That would involve fitting models to combinations of polio case data and ES surveillance results. Models that incorporate the waning patterns examined in this paper could be fit. Various approaches to fitting in the POMP package (King et al., 2016) could be used. Such fitting requires models that are simpler than Kid Risk models but that capture more realistic detail of where cases are occurring and where ES samples are being examined than in the models examined in this paper.

As information is gained on the likelihood of different waning scenarios, that information could then be used to indicate where location of ES sites will be most helpful for guiding decisions on when to stop the use of OPV. Currently sites are being located where the most problems in eliminating polio cases have been experienced. But one implication of adults sustaining prolonged low-level silent circulation as suggested by the analysis in this paper is that populations not exposed to OPV from kids might have more waned immunity and thus be more likely to sustain reinfection transmission chains given that the transmission conditions are right. In that case the location of ES might be chosen on the basis of how informative ES will be for assessing the overall risk of silent circulation rather than just to maximize the chances of a detection.

## Acknowledgements

This work was funded by two NIH MIDAS grants (U01GM110712 and 5U54GM111274) and a WHO grant 353558 TSA 2014/485861-0

## References

Abbink, F., Buisman, A.M., Doornbos, G., Woldman, J., Kimman, T.G., Conyn-van Spaendonck, M.A., 2005. Poliovirus-specific memory immunity in seronegative elderly people does not protect against virus excretion. J Infect Dis 191, 990–999.

Arie, S., 2014. Polio virus spreads from Syria to Iraq. BMJ 348, g2481.

Aylward, R.B., Alwan, A., 2014. Polio in syria. Lancet 383, 489–491.

Behrend, M.R., Hu, H., Nigmatulina, K.R., Eckhoff, P., 2014. A quantitative survey of the literature on poliovirus infection and immunity. Int J Infect Dis 18, 4–13.

Debanne, S.M., Rowland, D.Y., 1998. Statistical certification of eradication of poliomyelitis in the Americas. Math Biosci 150, 83–103.

Duintjer Tebbens, R.J., Kalkowska, D.A., Wassilak, S.G., Pallansch, M.A., Cochi, S.L., Thompson, K.M., 2014. The potential impact of expanding target age groups for polio immunization campaigns. BMC Infect Dis 14, 45.

Duintjer Tebbens, R.J., Pallansch, M.A., Chumakov, K.M., Halsey, N.A., Hovi, T., Minor, P.D., Modlin, J.F., Patriarca, P.A., Sutter, R.W., Wright, P.F., Wassilak, S.G., Cochi, S.L., Kim, J.H., Thompson, K.M., 2013a. Expert review on poliovirus immunity and transmission. Risk Anal 33, 544–605.

Duintjer Tebbens, R.J., Pallansch, M.A., Chumakov, K.M., Halsey, N.A., Hovi, T., Minor, P.D., Modlin, J.F., Patriarca, P.A., Sutter, R.W., Wright, P.F., Wassilak, S.G., Cochi, S.L., Kim, J.H., Thompson, K.M., 2013b. Review and assessment of poliovirus immunity and transmission: synthesis of knowledge gaps and identification of research needs. Risk Anal 33, 606–646.

Duintjer Tebbens, R.J., Pallansch, M.A., Kalkowska, D.A., Wassilak, S.G., Cochi, S.L., Thompson, K.M., 2013c. Characterizing poliovirus transmission and evolution: insights from modeling experiences with wild and vaccine-related polioviruses. Risk Anal 33, 703–749.

Duintjer Tebbens, R.J., Pallansch, M.A., Kim, J.H., Burns, C.C., Kew, O.M., Oberste, M.S., Diop, O.M., Wassilak, S.G., Cochi, S.L., Thompson, K.M., 2013d. Oral poliovirus vaccine evolution and insights relevant to modeling the risks of circulating vaccine-derived polioviruses (cVDPVs). Risk Anal 33, 680–702.

Duintjer Tebbens, R.J., Pallansch, M.A., Wassilak, S.G., Cochi, S.L., Thompson, K.M., 2015. Combinations of Quality and Frequency of Immunization Activities to Stop and Prevent Poliovirus Transmission in the High-Risk Area of Northwest Nigeria. PLoS One 10, e0130123.

Duintjer Tebbens, R.J., Thompson, K.M., 2014. Modeling the potential role of inactivated poliovirus vaccine to manage the risks of oral poliovirus vaccine cessation. J Infect Dis 210 Suppl 1, S485–497.

Duintjer Tebbens, R.J., Thompson, K.M., 2015. Managing the risk of circulating vaccine-derived poliovirus during the endgame: oral poliovirus vaccine needs. BMC Infect Dis 15, 390.

Eichner, M., Dietz, K., 1996. Eradication of poliomyelitis: when can one be sure that polio virus transmission has been terminated? Am J Epidemiol 143, 816–822.

Famulare, M., 2015. Has Wild Poliovirus Been Eliminated from Nigeria? PLoS One 10, e0135765.

Famulare, M., Chang, S., Iber, J., Zhao, K., Adeniji, J.A., Bukbuk, D., Baba, M., Behrend, M., Burns, C.C., Oberste, M.S., 2015. Sabin vaccine reversion in the field: a comprehensive analysis of Sabin-like poliovirus isolates in Nigeria. J Virol.

Fine, P.E., Carneiro, I.A., 1999. Transmissibility and persistence of oral polio vaccine viruses: implications for the global poliomyelitis eradication initiative. Am J Epidemiol 150, 1001–1021.

GPEI, 2015. POLIO ENVIRONMENTAL SURVEILLANCE EXPANSION PLAN.

GPEI, 2016. Global Polio Eradication Initiative: Data and Monitoring: Nigeria, p. http://www.polioeradication.org/dataandmonitoring/poliothisweek.aspx.

Grassly, N.C., 2013. The final stages of the global eradication of poliomyelitis. Philos Trans R Soc Lond B Biol Sci 368, 20120140.

Hagan, J.E., Wassilak, S.G., Craig, A.S., Tangermann, R.H., Diop, O.M., Burns, C.C., Quddus, A., 2015. Progress toward polio eradication - worldwide, 2014–2015. MMWR Morb Mortal Wkly Rep 64, 527–531.

Jafari, H., Deshpande, J.M., Sutter, R.W., Bahl, S., Verma, H., Ahmad, M., Kunwar, A., Vishwakarma, R., Agarwal, A., Jain, S., Estivariz, C., Sethi, R., Molodecky, N.A., Grassly, N.C., Pallansch, M.A., Chatterjee, A., Aylward, R.B., 2014. Polio eradication. Efficacy of inactivated poliovirus vaccine in India. Science 345, 922–925.

Kalkowska, D.A., Duintjer Tebbens, R.J., Pallansch, M.A., Cochi, S.L., Wassilak, S.G., Thompson, K.M., 2015. Modeling undetected live poliovirus circulation after apparent interruption of transmission: implications for surveillance and vaccination. BMC Infect Dis 15, 66.

Kalkowska, D.A., Duintjer Tebbens, R.J., Thompson, K.M., 2012. The probability of undetected wild poliovirus circulation after apparent global interruption of transmission. Am J Epidemiol 175, 936–949.

King, A.A., Ionides, E., Breto, C., Ellner, S.P., Ferrari, M.J., Kendall, B.E., Lavine, M., Nguyen, D., Reuman, D.C., Wearing, H., Wood, S.N., Funk, S., 2016. Package “pomp”.

Manor, Y., Shulman, L.M., Kaliner, E., Hindiyeh, M., Ram, D., Sofer, D., Moran-Gilad, J., Lev, B., Grotto, I., Gamzu, R., Mendelson, E., 2014. Intensified environmental surveillance supporting the response to wild poliovirus type 1 silent circulation in Israel, 2013. Euro Surveill 19, 20708.

Mayer, B.T., Eisenberg, J.N., Henry, C.J., Gomes, M.G., Ionides, E.L., Koopman, J.S., 2013. Successes and shortcomings of polio eradication: a transmission modeling analysis. American Journal of Epidemiology 177, 1236–1245.

Nosek, B.A., Alter, G., Banks, G.C., Borsboom, D., Bowman, S.D., Breckler, S.J., Buck, S., Chambers, C.D., Chin, G., Christensen, G., Contestabile, M., Dafoe, A., Eich, E., Freese, J., Glennerster, R., Goroff, D., Green, D.P., Hesse, B., Humphreys, M., Ishiyama, J., Karlan, D., Kraut, A., Lupia, A., Mabry, P., Madon, T.A., Malhotra, N., Mayo-Wilson, E., McNutt, M., Miguel, E., Paluck, E.L., Simonsohn, U., Soderberg, C., Spellman, B.A., Turitto, J., VandenBos, G., Vazire, S., Wagenmakers, E.J., Wilson, R., Yarkoni, T., 2015. SCIENTIFIC STANDARDS. Promoting an open research culture. Science 348, 1422–1425.

Oster, G., Macey, R., 2015. Berkeley Madonna, Modeling and analysis of dynamic systems, Berkeley, California, p. Main web page for purchase and for getting documentation.

Porter, K.A., Diop, O.M., Burns, C.C., Tangermann, R.H., Wassilak, S.G., 2015. Tracking progress toward polio eradication - worldwide, 2013–2014. MMWR Morb Mortal Wkly Rep 64, 415–420.

Pujol, J.M., Eisenberg, J.E., Haas, C.N., Koopman, J.S., 2009. The Effect of Ongoing Exposure Dynamics in Dose Response Relationships. PLoS Comput Biol 5, e1000399.

Shulman, L.M., Gavrilin, E., Jorba, J., Martin, J., Burns, C.C., Manor, Y., Moran-Gilad, J., Sofer, D., Hindiyeh, M.Y., Gamzu, R., Mendelson, E., Grotto, I., 2014a. Molecular epidemiology of silent introduction and sustained transmission of wild poliovirus type 1, Israel, 2013. Euro Surveill 19, 20709.

Shulman, L.M., Martin, J., Sofer, D., Burns, C.C., Manor, Y., Hindiyeh, M., Gavrilin, E., Wilton, T., Moran-Gilad, J., Gamzo, R., Mendelson, E., Grotto, I., 2014b. Genetic Analysis and Characterization of Wild Poliovirus Type 1 During Sustained Transmission in a Population With >95% Vaccine Coverage, Israel 2013. Clin Infect Dis.

Shulman, L.M., Mendelson, E., Anis, E., Bassal, R., Gdalevich, M., Hindiyeh, M., Kaliner, E., Kopel, E., Manor, Y., Moran-Gilad, J., Ram, D., Sofer, D., Somekh, E., Tasher, D., Weil, M., Gamzu, R., Grotto, I., 2014c. Laboratory challenges in response to silent introduction and sustained transmission of wild poliovirus type 1 in Israel during 2013. J Infect Dis 210 Suppl 1, S304–314.

Teunis, P.F.M., van Eijkeren, J.C.H., de Graafd, W.F., Bonacic Marinovic, A., Kretzschmar, M.E.E., 2016. Linking the seroresponse to infection to within-host heterogeneity inantibody production. Epidemics 16, 33–39.

Thompson, K.M., 2013. Modeling poliovirus risks and the legacy of polio eradication. Risk Anal 33, 505–515.

Thompson, K.M., Duintjer Tebbens, R.J., 2014a. Modeling the dynamics of oral poliovirus vaccine cessation. J Infect Dis 210 Suppl 1, S475–484.

Thompson, K.M., Duintjer Tebbens, R.J., 2014b. National choices related to inactivated poliovirus vaccine, innovation and the endgame of global polio eradication. Expert Rev Vaccines 13, 221–234.

Thompson, K.M., Kalkowska, D.A., Duintjer Tebbens, R.J., 2015a. Managing population immunity to reduce or eliminate the risks of circulation following the importation of polioviruses. Vaccine.

Thompson, K.M., Kalkowska, D.A., Duintjer Tebbens, R.J., 2015b. Managing population immunity to reduce or eliminate the risks of circulation following the importation of polioviruses. Vaccine 33, 1568–1577.

Thompson, K.M., Pallansch, M.A., Duintjer Tebbens, R.J., Wassilak, S.G., Kim, J.H., Cochi, S.L., 2013a. Preeradication vaccine policy options for poliovirus infection and disease control. Risk Anal 33, 516–543.

Thompson, K.M., Pallansch, M.A., Tebbens, R.J., Wassilak, S.G., Cochi, S.L., 2013b. Modeling population immunity to support efforts to end the transmission of live polioviruses. Risk Anal 33, 647–663.

Thompson, K.M., Strebel, P.M., Dabbagh, A., Cherian, T., Cochi, S.L., 2013c. Enabling implementation of the Global Vaccine Action Plan: developing investment cases to achieve targets for measles and rubella prevention. Vaccine 31 Suppl 2, B149–156.

Thompson, K.M., Tebbens, R.J., 2012. Current polio global eradication and control policy options: perspectives from modeling and prerequisites for oral poliovirus vaccine cessation. Expert Rev Vaccines 11, 449–459.

Volz, E.M., Ionides, E., Romero-Severson, E.O., Brandt, M.G., Mokotoff, E., Koopman, J.S., 2013. HIV-1 transmission during early infection in men who have sex with men: a phylodynamic analysis. PLoS Med 10, e1001568; discussion e1001568.

Wagner, B.G., Behrend, M.R., Klein, D.J., Upfill-Brown, A.M., Eckhoff, P.A., Hu, H., 2014. Quantifying the impact of expanded age group campaigns for polio eradication. PLoS One 9, e113538.

WHO, 2015. Polio Eradication and Endgame Midterm Review 2015, in: WHO (Ed.), WHO/Polio/15.04. WHO, Geneva.

